# Application of Long Read Sequencing to Determine Expressed Antigen Diversity in *Trypanosoma Brucei* Infections

**DOI:** 10.1101/432245

**Authors:** Siddharth Jayaraman, Claire Harris, Edith Paxton, Anne-Marie Donachie, Heli Vaikkinen, Richard McCulloch, James P. J. Hall, John Kenny, Luca Lenzi, Christiane Hertz-Fowler, Christina Cobbold, Richard Reeve, Tom Michoel, Liam J. Morrison

**Affiliations:** Roslin Institute, University of Edinburgh, Easter Bush, Midlothian, EH25 9RG, United Kingdom; Boyd Orr Centre for Population and Ecosystem Health, Institute of Biodiversity, Animal Health and Comparative Medicine, College of Medical and Life Sciences, University of Glasgow, 120 University Place, Glasgow G12 8TA, United Kingdom; Wellcome Centre for Molecular Parasitology, Institute of Infection, Immunity and Inflammation, College of Medical and Life Sciences, University of Glasgow, 120 University Place, Glasgow G12 8TA, United Kingdom; Department of Biology, Wentworth Way, University of York, York YO10 5DD, United Kingdom; Centre for Genomic Research, Institute of Integrative Biology, University of Liverpool, Crown Street, Liverpool L69 7ZB, United Kingdom; School of Mathematics and Statistics, University of Glasgow, University Place, Glasgow, G12 8QS, United Kingdom

**Author notes:** 00441316519470. LM and TM are joint last authors.

## Abstract

Antigenic variation is employed by many pathogens to evade the host immune response, and *Trypanosoma brucei* has evolved a complex system to achieve this phenotype, involving sequential use of variant surface glycoprotein (VSG) genes encoded from a large repertoire of ∼2,000 alleles. *T. brucei* express multiple, sometimes closely related, VSGs in a population at any one time, and the ability to resolve and analyse this diversity has been limited. We applied long read sequencing (PacBio) to VSG amplicons generated from blood extracted from batches of mice sacrificed at time points (days 3, 6, 10 and 12) post-infection with T. brucei TREU927. The data showed that long read sequencing is reliable for resolving allelic differences between VSGs, and demonstrated that there is significant expressed diversity (449 VSGs detected across 20 mice) and across the timeframe of study there was a clear semi-reproducible pattern of expressed diversity (median of 27 VSGs per sample at day 3 post infection (p.i.), 82 VSGs at day 6 p.i., 187 VSGs at day 10 p.i. and 132 VSGs by day 12 p.i.). There was also consistent detection of one VSG dominating expression across replicates at days 3 and 6, and emergence of a second dominant VSG across replicates by day 12. The innovative application of ecological diversity analysis to VSG reads enabled characterisation of hierarchical VSG expression in the dataset, and resulted in a novel method for analysing such patterns of variation. Additionally, the long read approach allowed detection of mosaic VSG expression from very few reads – this was observed as early as day 3, the earliest that such events have been detected. Therefore, our results indicate that long read analysis is a reliable tool for resolving diverse allele expression profiles, and provides novel insights into the complexity and nature of VSG expression in trypanosomes, revealing significantly higher diversity than previously shown and identifying mosaic gene formation unprecedentedly early during the infection process.

## Introduction

Antigenic variation is used by many pathogens as a means of staying one step ahead of the host’s adaptive immune response. *Trypanosoma brucei* has long been a paradigm for the study of antigenic variation, and the protein responsible, the variable surface glycoprotein (VSG) has been the focus of much research [1–3]. Each trypanosome in a population expresses a single species of protein, and an inherent, parasite-driven switching process causes a proportion of the population to replace their active VSG gene with a different VSG gene, resulting in the expression of a protein in those cells with different epitopes exposed to the host immune system (at a rate of up to 10^−2^ switches per cell/generation [4]). The post-genomic era has revealed *T. brucei*’s antigenic variation system to be unrivalled in its elaboration, particularly in terms of the scale of the numbers of genes that comprise the VSG family. Sequencing the genome of *T. brucei* has uncovered a gene family much greater in numbers and complexity than was previously thought. Characterisation to date suggests that at least 2,000 VSG genes are in the genome of each trypanosome, providing a spectacularly large repertoire of potential antigens [5, 6], particularly when compared to other pathogens that undergo antigenic variation, such as *Plasmodium Falciparum* (60 genes in PfEMP1 family [7]), *Anaplasma marginale* (∼10 members in the *msp2* & *msp3* gene families [8]), and *Borrelia burgdorferi* (15 members in the *vls* gene family[9]).

The scale of the gene family size is also reflected in the complexity of switching mechanisms employed to change the identity of the surface antigen. The VSGs are expressed from one of approximately 20 bloodstream expression sites (BES)[10], the active expression occurring in a dedicated sub-nuclear organelle, the expression site body (ESB)[11], with the remainder of BESs being transcriptionally silent. A minor mechanism of VSG switching, accounting for only approximately 10% of events in wild type trypanosomes [12], is to turn off the transcription of the active BES and activate one of the silent BESs (‘transcriptional switching’). However, the majority of switching is through replacing the gene sequence in the active BES via gene duplication, which involves the copying of variable amounts of sequence, ranging from within the gene to the whole telomere [13, 14]. Insights into mechanisms involved in switching suggest that replacing allelic sequence is driven by DNA recombination, and DNA repair/homologous recombination pathways and proteins (e.g. RAD51) have been identified to be involved in the gene duplication process of VSG switching [15] (reviewed in [16]). A further layer of complexity is the construction of novel VSG sequences in the BES from multiple donor VSG sequences, a form of segmental gene conversion termed ‘mosaic’ gene conversion [17, 18]. Mosaic gene formation was previously considered to be a rare and minor mechanistic component of overall VSG switching in an infection (e.g. [14]). However, the revelation upon the sequencing of the *T. brucei* genome that a significant proportion of the VSG repertoire (80–90%) consisted of pseudogenes [19] that cannot be expressed as functional proteins began to alter that perception [5, 20]. It has become clear from subsequent experimental work that mosaic gene formation is an integral component of VSG switching, particularly after the early stages of infection (i.e. beyond the first peak of parasitaemia in mouse infections)[5, 21].

One of the challenges of analysing VSG expression in vivo, and in particular gaining an accurate measurement of the level of expressed diversity given the extent of the VSG repertoire (i.e. to what extent is the repertoire actually used during infection), has been the relatively limited resolution of available techniques – in particular the manual cloning and sequencing of individual VSG cDNAs that has been undertaken in recent studies [5, 21]). While this approach clearly provides accurate data at the level of individual VSG transcripts, the limitations have undoubtedly resulted in a low estimate of the diversity and complexity of VSG expression at the population level, and particularly with respect to minor variant populations. Additionally, although transcriptomics potentially provides the ability to overcome the resolution limitations of manually cloning and sequencing transcripts, the application of RNAseq to VSG expression from *in vivo* samples has long been deemed challenging, due to the requirement for assembling multiple closely related alleles from a mixed population using short reads of 100–200 base pairs (e.g. Illumina) – this has similarly been an issue when attempting to resolve, for example, the diversity of the mammalian B cell repertoire underpinning the antibody response (e.g. [22]). However, a recent study subjected in vivo samples to Illumina sequencing (100bp, single-end reads) and demonstrated the utility of transcriptomics in terms of increased resolution [23], and were able to detect minor populations (0.1% of population) and up to 79 variants at a time point, although they were not able to identify significant mosaic gene expression.

Long read sequencing potentially provides the ability to further increase our resolution, particularly as the length of reads commonly reached with such technologies (average read length in Pacbio, for example, is quoted as 10–20,000 bp; http://www.pacb.com/smrt-science/smrt-sequencing/read-lengths/) far exceeds the length of the VSG transcript (approximately 1600 bp), meaning that the issue of assembly of closely VSGs from multiple reads should be bypassed. Here, we present analysis of VSG expression from replicate in vivo *T. brucei* TREU927 infections in mice at 4 time points over 12 days using almost 500,000 Pacbio Sequencing reads. We demonstrate that long read technologies provide significant advantages for analysing the diversity of VSG expression. Our data suggest that the VSG population comprises significantly more variants even at an early stage of infection (up to 190 variants at day 10 post-infection), that the pattern of VSG expression is surprisingly reproducible (using the novel application of ecological diversity indices), and that mosaic gene expression can be detected much earlier in infection than has been possible previously. Our data also provides insights into the nature of mutations introduced by Pacbio long-read sequencing technology, as the dataset includes significant coverage of one sequence (>140,000 reads).

## Results

### Long read sequencing maps the VSG transcriptome at unprecedented resolution

Using PacBio long read RNA sequencing of 20 blood samples enriched for VSG transcripts from replicate in vivo *T. brucei* TREU927 infections in mice at 3, 6, 10 and 12 days post infection, we obtained 486,343 reads with an average read length of insert of 1,569 bp (Table 1, Figure 1B). Reads were filtered by length (1400–2000bp) based upon both literature on VSG genes [21, 24] and the read distribution in our dataset (Figure 1B) to remove reads resulting from sequencing artefacts and shorter fragments (i.e. partial reads), and on the basis of similarity to known VSGs (blastn ≥60% alignment against TriTrypDB-v26 [25]) (Figure 1C). This resulted in a dataset of 296,937 reads, with an average of 6.50 full passes per read (summarised in Table 1; full data in S1 Table), and a robust identification of a donor gene for the N-Terminal domain (NTD), where we have high confidence in the reads containing all of the features necessary to be consistent with being full length VSG transcripts. The 296,937 reads represent a total of 449 VSGs (74.77% of VSG a-type and 25.22% VSG b-type [24]) across 20 samples, with the number of reads per VSG following a power-law distribution (Figure 1D), and provide a unique insight into the in vivo VSG transcriptome across time and animal replicate.

**Figure 1.**
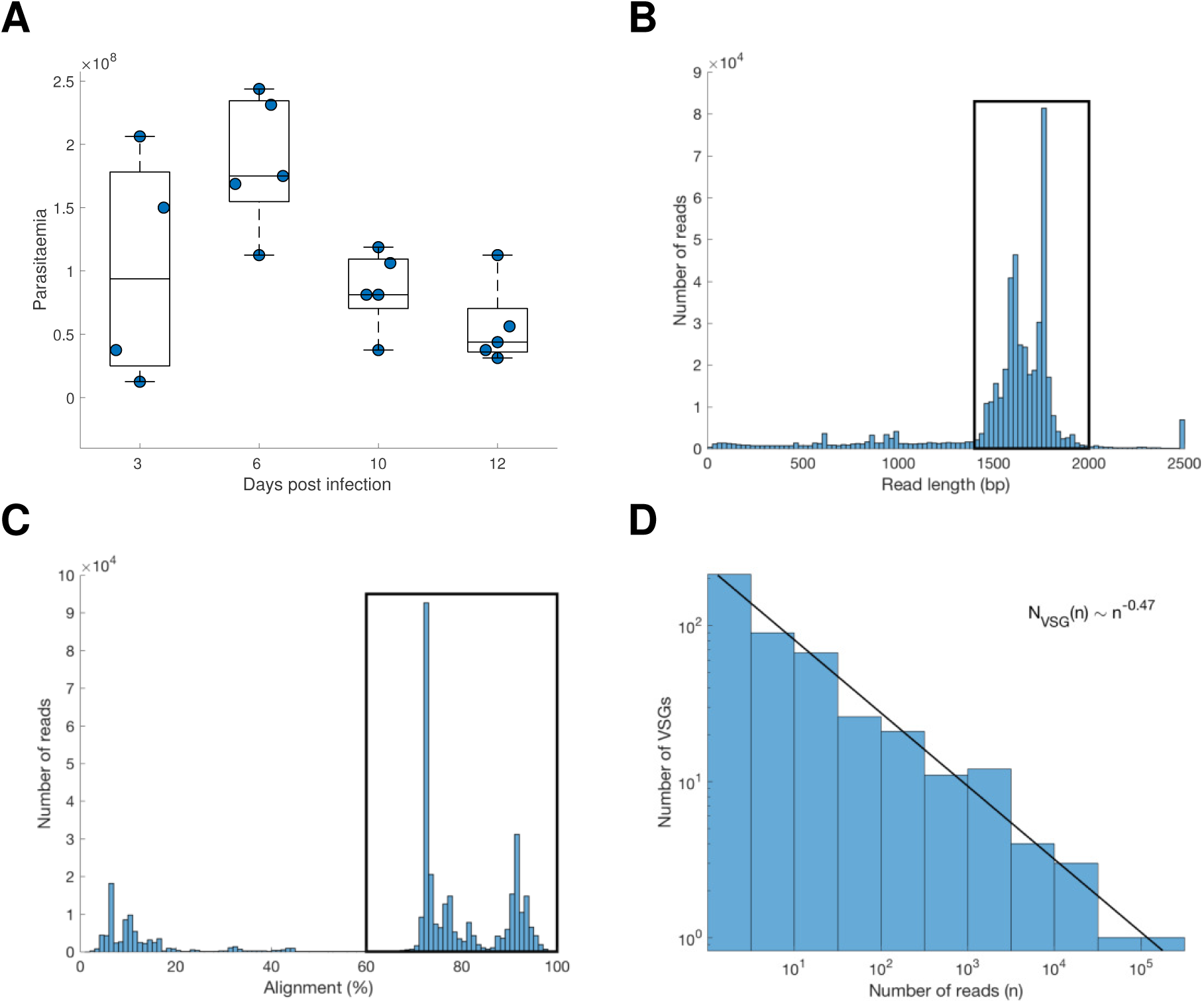
(A) Parasitaemia measurements for the batches of mice sacrificed at days 3, 6, 9 and 12 days post-infection. A summary of analysis of pacbio data generated from these samples is shown, including main decision steps (black boxes represent data filtering steps; data within black boxes was included), (B) read length distribution, (C) alignment of reads to VSG, (D) correlation of read number with number of VSGs identified.

**Table 1.**
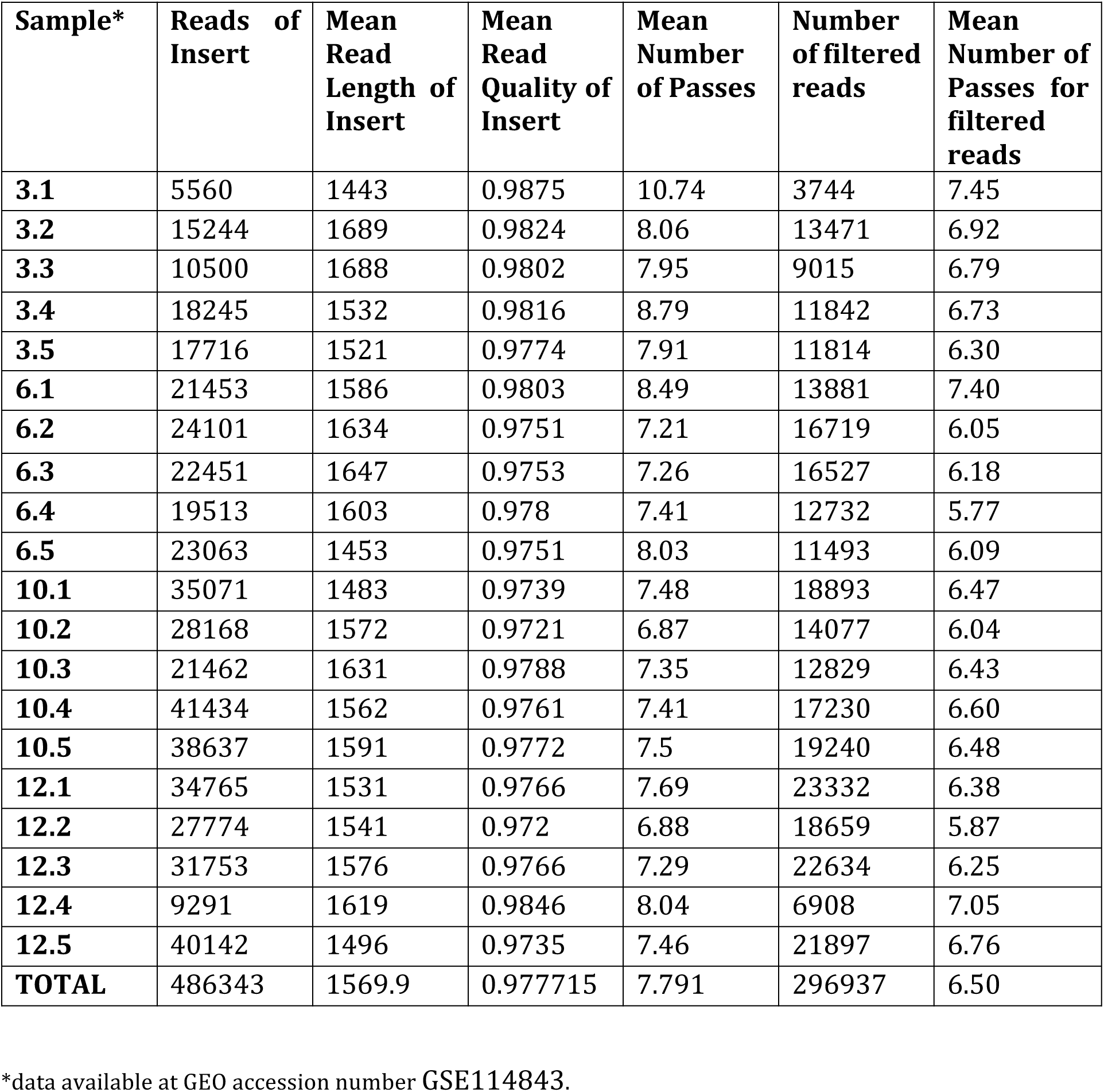
Summary of data per sample.

### PacBio sequences contain random errors even at high number of full passes

ORFs were identified in the 296,937 reads with a conservative minimum nucleotide size of 1200 nucleotides (reported size ranges of VSG NTDs and C-Terminal domains [CTDs] are approximately 300–350 and 100 amino acids, respectively [5, 21, 24]). Surprisingly, only 33,234 reads (11%) resulted in predicted ORFs. Although the percentage of reads with predicted ORF increased with increasing number of full passes, it remained well below 50% even for reads having 10 full passes or more (Figure 2A). Since the distribution of the number of reads with a detectable ORF over all VSGs was similar to total expression level distribution (Table 2), we hypothesize that the lack of identified ORFs was due to random sequencing errors rather than any systematic biases in the data, despite PacBio claiming an accuracy of more than 99% for reads with 15-fold coverage [26]. To investigate this hypothesis in more detail, we focused on the most abundant VSG (Tb08.27P2.380, 1551bp, 141,822 high-confidence reads) and annotated each discrepant base pair of each aligned read as either an insertion, deletion or mismatch with respect to the Tb08.27P2.380 reference genome sequence. All reads had an alignment score greater than 90% over the first 1266bp (the N-Terminal domain) (Figure 2B). The distribution of sequence errors showed a clear bimodal pattern across the N-Terminal domain, with 145 nucleotide positions having a consistent mismatch (131), deletion (10) or insertion (4) across more than 80% of the reads, and 1,112 nucleotide positions having errors in at least one but fewer than 10% of reads (Figure 2B). This suggests that the former represent genuine mutations already present in our inoculum (with respect to the reference genome sequence), whereas the latter represent either random sequencing errors introduced by Pacbio or low level genuine mutations that we cannot currently distinguish from Pacbio error. Previous studies have indicated accumulation of mutations over time in expressed VSGs, and we examined this in our data for reads aligning to Tb08.27P2.380 (for the N-terminal domain) by assessing the error rate for mismatches, insertions and deletions (S2 Figure). While these data indicated statistical support for differences in the data distribution across time points for all 3 mutation classes, due to the skewed nature of the data distribution (most bases have an error rate close to zero) this conclusion must be treated with a degree of caution. The assertion that the errors present in > 80% of reads were ‘genuine’ mutations was further supported by these 145 mutations being consistently present in PCR amplicons generated from cDNA extracted from multiple samples that were sequenced by Sanger sequencing (n = 12 for Tb08.27P2.380; representing sequences independently cloned and sequenced from 7 mice on days 3, 6 and 10; data not shown). Deletions were by far the most common Pacbio-introduced error (average per-base error rate of 3.2% across the N-terminal domain sequence), followed by mismatches (0.8%) and insertions (0.3%) (Figure 2C), in agreement with what has been reported before [27]. Consistent with the ORF prediction pattern (Figure 2A), the overall error percentage was lower for reads with higher number of passes, but introduced sequencing errors (i.e. interpreted as mutations not present in the genome of the inoculated trypanosomes) remained present at more than 1000 nucleotide positions even for reads with 10 passes (Figure 2D). The nature of our data, comprising > 141,000 reads of the same sequence, therefore provides an unusually robust insight into the nature of Pacbio errors and the caveats that must be placed upon interpretation of such data, as most studies involve much less coverage per single base pair.

**Figure 2.**
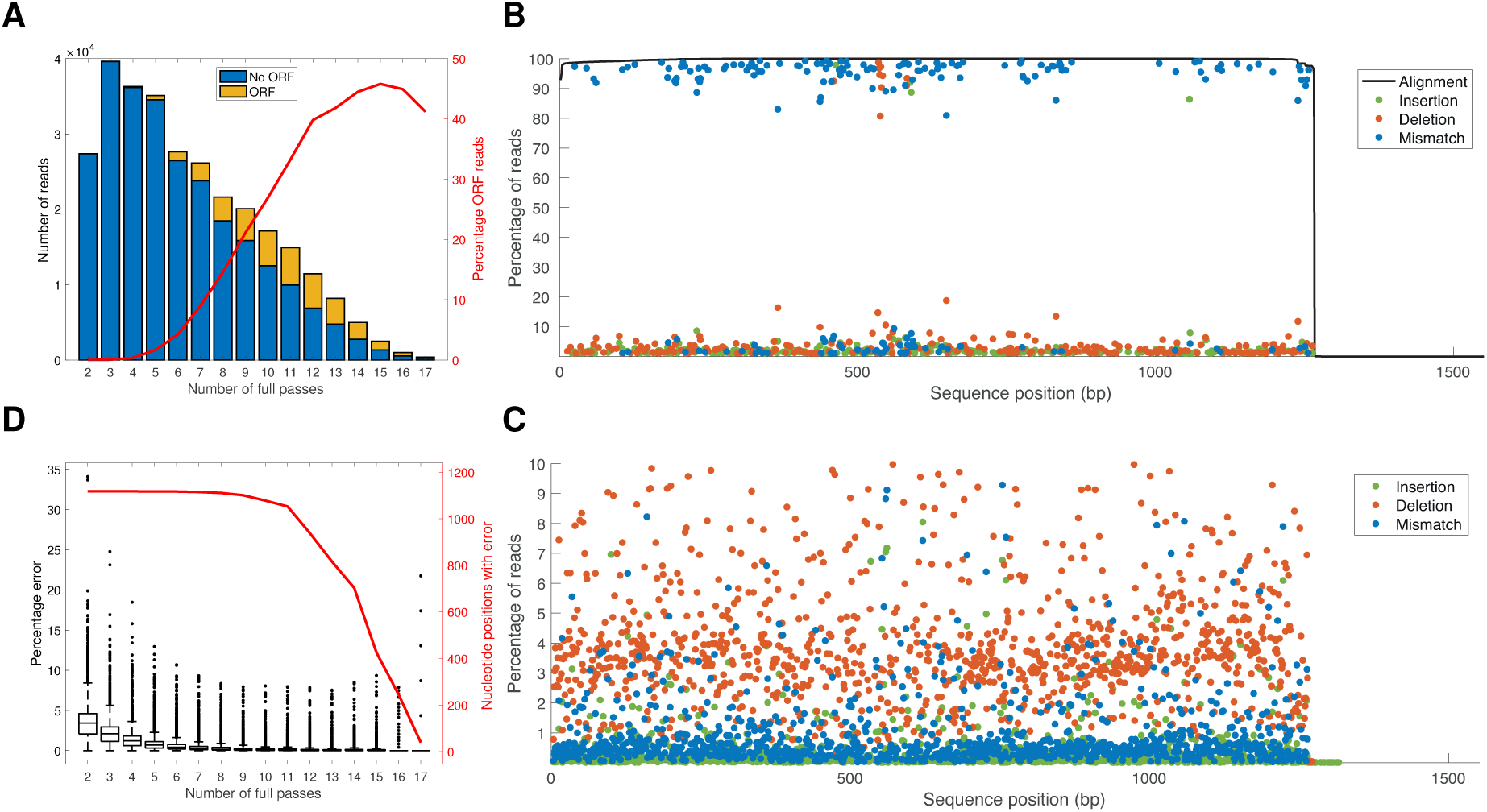
Analysis of pacbio sequence data: (A) number of reads aligning to VSGs per number of full passes, with proportion of those reads comprising predicted ORF in yellow; red line shows percentage of reads at each threshold of full pass number that contained a predicted VSG ORF. (B) percentage of reads at each position in the N-Terminal domain that contained a mutation with respect to the reference genome sequence for VSG Tb08.27P2.380; alignment coverage is shown by the black line, insertions by green dot, deletions by red dot and mismatch by blue dot. Highlighted dots (with circle around dot) are those that were different to both the reference sequence and Sanger sequence cloned VSG sequences from the inoculated stabilate at a high percentage of reads at that position (>70%), (C) focussed representation of data in 2B, with only mutations with respect to the genome reference sequence <10% at each position in the N-Terminal domain VSG Tb08.27P2.380 shown; insertions shown by green dot, deletions by red dot and mismatch by blue dot. (D) percentage error rate plotted against number of full passes; red line indicates number of nucleotide positions for those aligning to VSG Tb08.27P2.380 that contained an error with respect to genome reference sequence against number of full passes.

**Table 2.** Top 20 variants; number of reads across dataset as measured by (i) mapping to reference VSG database and by (ii) sequence clustering; the relevant TREU927 reference VSG is indicated in the first column.

| **VSG reference gene** | **Reads by mapping to VSGs** | **Percentage of total** | **Reads by sequence clustering\*** | **Percentage of total** | **Hall Sets** |
| --- | --- | --- | --- | --- | --- |
| Total | **296937** | 100 | 33205 | 100 |  |
| Tb08.27P2.380 | 141822 | 47.76 | 14543 | 43.80 | Set_23 |
| Tb09.v4.0077 | 46264 | 15.58 | 5951 | 17.92 | Set_08 |
| Tb927.4.5730 | 23643 | 7.96 | 2499 | 7.53 |  |
| Tb927.10.10 | 13167 | 4.43 | 1294 | 3.90 |  |
| Tb11.v5.0932 | 10187 | 3.43 | 1435 | 4.32 |  |
| Tb927.9.300 | 8751 | 2.94 | 913 | 2.75 |  |
| Tb09.v4.0088 | 7947 | 2.67 | 1223 | 3.68 | Set_36 |
| Tb05.5K5.330 | 7865 | 2.64 | 874 | 2.63 | Set_22 |
| Tb927.9.16490 | 3669 | 1.23 | 413 | 1.24 |  |
| Tb927.3.480 | 2984 | 1 | 368 | 1.11 |  |
| Tb11.57.0047 | 2783 | 0.93 | 379 | 1.14 |  |
| Tb927.11.20300 | 2353 | 0.79 | 247 | 0.74 |  |
| Tb927.1.05 | 1990 | 0.67 | 280 | 0.84 |  |
| Tb927.4.5570 | 1622 | 0.54 | 200 | 0.60 |  |
| Tb10.v4.0061 | 1414 | 0.47 | 181 | 0.55 | Set_04 |
| Tb11.v5.0599 | 1400 | 0.47 | - | - |  |
| Tb927.9.580 | 1276 | 0.42 | - | - |  |
| Tb927.5.4700 | 1168 | 0.39 | 175 | 0.53 |  |
| Tb09.v4.0075 | 1133 | 0.38 | 140 | 0.42 | Set_35 |
| Tb11.1451 | 1126 | 0.37 | 121 | 0.36 |  |

\* The subset of reads with predicted ORFs was used for the clustering algorithm analysis.

### VSG population comprises more variants than expected and is highly reproducible

Our data demonstrate that we can detect multiple VSGs in each sample, and that we can identify changes in VSG expression and diversity over time. We identified a median of 27 unique VSGs per sample at day 3 post infection (p.i.), which progressed to 82 VSGs at day 6 p.i., peaking at 187 VSGs at day 10 p.i. and reducing to 132 VSGs by day 12 p.i. (Figure 3A). When identified VSGs that mapped to single reads from single samples were removed, this resulted in an identification of 334 VSGs (median of 27, 81, 170 and 126 VSGs per sample at 3, 6, 10, and 12 days p.i., respectively).

**Figure 3.**
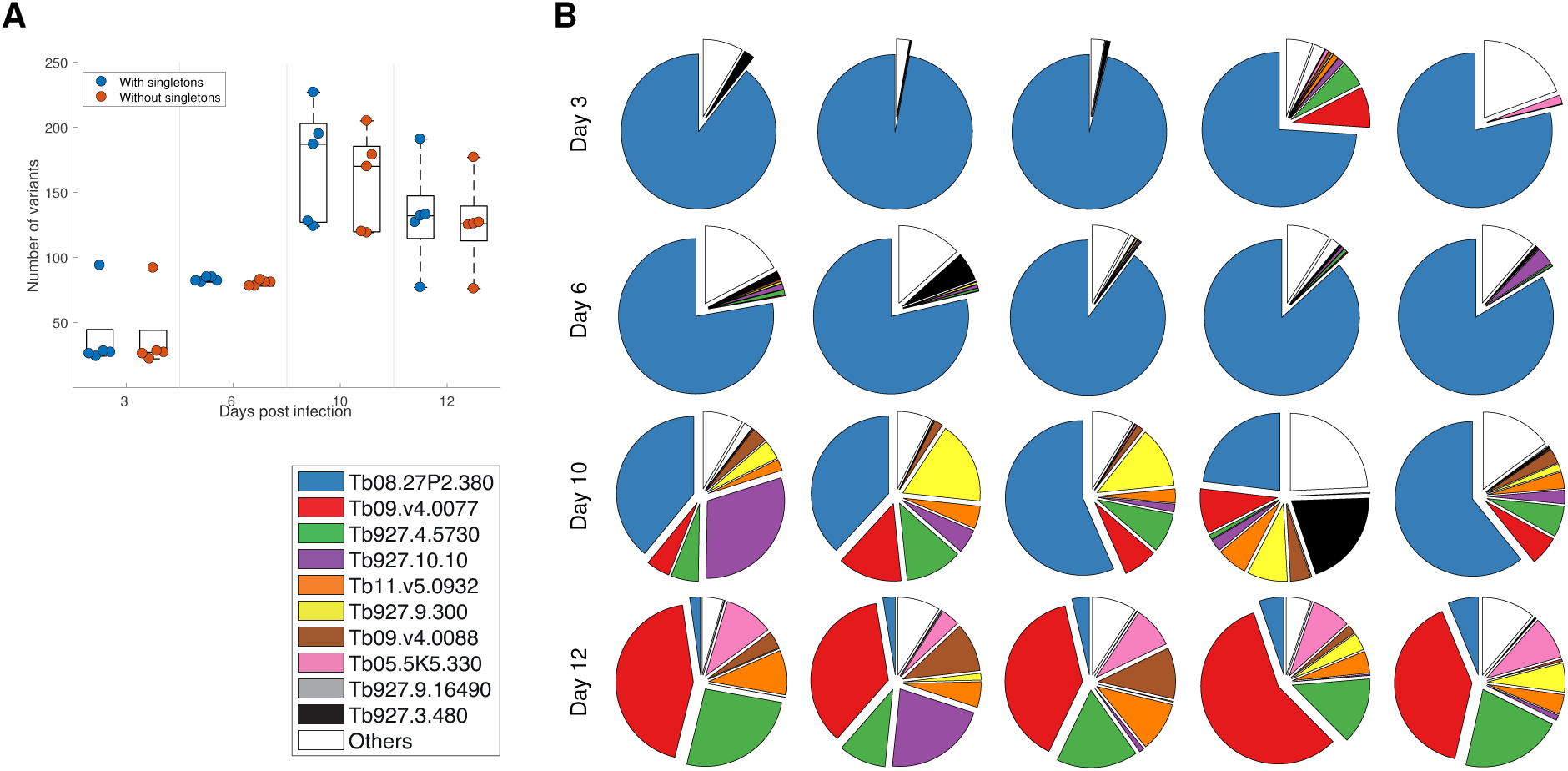
VSGs per sample. (A) Average number of donor VSGs mapped at each time point, plotted with and without VSGs identified from single reads of at least one full pass (‘singletons’), (B) Proportion of reads that map to identified donor VSGs for each mouse and each timepoint; reads that map to the 10 most abundant identified donors (across the whole dataset from 20 mice) are shown; reads that map to donor genes other than these ten are represented as ‘others’).

Not only were the number of distinct VSGs consistent across samples for the same time point, but the expression pattern (proportion of reads per sample mapping to particular VSGs) was also highly reproducible between samples and over time (Figure 3B), albeit bearing in mind that these analyses are of batches of mice at four different time points rather than longitudinal samples of the same mice. The VSG that is dominant at day 3 (Tb08.27P2.380), presumably introduced as the dominant VSG in the inoculum, remains dominant in all mice at day 6, but is only the VSG with the most reads aligned in only two of five mice at day 10. Interestingly, by day 12, the VSG with the most reads per sample is the same in all five mice (Tb09.v4.0077) and this VSG was also most common at similar timepoint in previous analyses [21]. Additionally, the other eight VSGs that reads map to in mice at days 10 and 12 (Tb927.4.5730, Tb927.10.10, Tb11.v5.0932, Tb927.9.300, Tb09.v4.0088, Tb05.5K5.330, Tb927.9.16490 and Tb927.3.480; Figure 3B) are present in all ten mice suggesting a degree of conservation in the sequential expression of VSGs in independent infections, consistent with previous observations [21, 28, 29]. However, in all mice there were reads that mapped to VSGs distinct to these most favoured 10 VSGs (‘others’ in Figure 3B, which account for 10.36% of all VSG-mapped reads), and in some mice this proportion was particularly high (e.g. mice 3.5 6.1 and 10.4; Figure 3B). This is particularly evident at day 6, where although the dominant VSG (Tb08.27P2.380) makes up most reads, the majority of reads that do not map to Tb08.27P2.380 map to VSGs other than the other top 9 VSGs in all mice. Additionally, at Day 10 we observe both the greatest number of VSGs and the least domination by any single VSG, but the proportion of ‘others’ either reduces or remains stable. These analyses combine to indicate that while there is a broad hierarchy in expression, with dominant VSGs at the beginning and end of infections, in between these timepoints there is a degree of stochasticity in the system – although eight VSGs comprise the majority of reads that do not map to either of the two dominant VSGs, the relative proportion of these ‘minority’ VSGs is not consistent, and there are furthermore many VSGs that are expressed at very low levels in all mice.

The analysis described thus far (Figure 3) has not taken into account any sequence similarity between VSGs, but relied on mapping reads to identified VSGs in the reference database. To analyse the population diversity of VSGs within and across samples using a method that is independent of mapping to existing databases (which are likely to be incomplete), we applied information theoretic measures more commonly used to quantify the biodiversity of ecosystems [30]. This approach initially applied a clustering algorithm to a proportion of reads (n = 33,205; comprising reads with predicted ORF) in order to enable identification of the reads that clustered on the basis of sequence similarity, as putatively distinct VSGs (Figure 4A). These data showed significant congruity with those described for the VSG mapping approach described above (Table 2). The top 10 clusters comprised 89.34% of all reads, compared to 89.68% for the VSG mapping approach, and the relative proportion of reads that either map to VSGs or cluster by sequence similarity is very comparable for the 10 most abundant VSGs (Table 2). These data indicate that the clustering algorithm applied was robust in terms of identifying individual VSGs, and therefore indicated a very similar pattern of a dominant early VSG, followed by an intermediate period of significant greater VSG diversity, ending up with a second dominant VSG by day 12.

**Figure 4.**
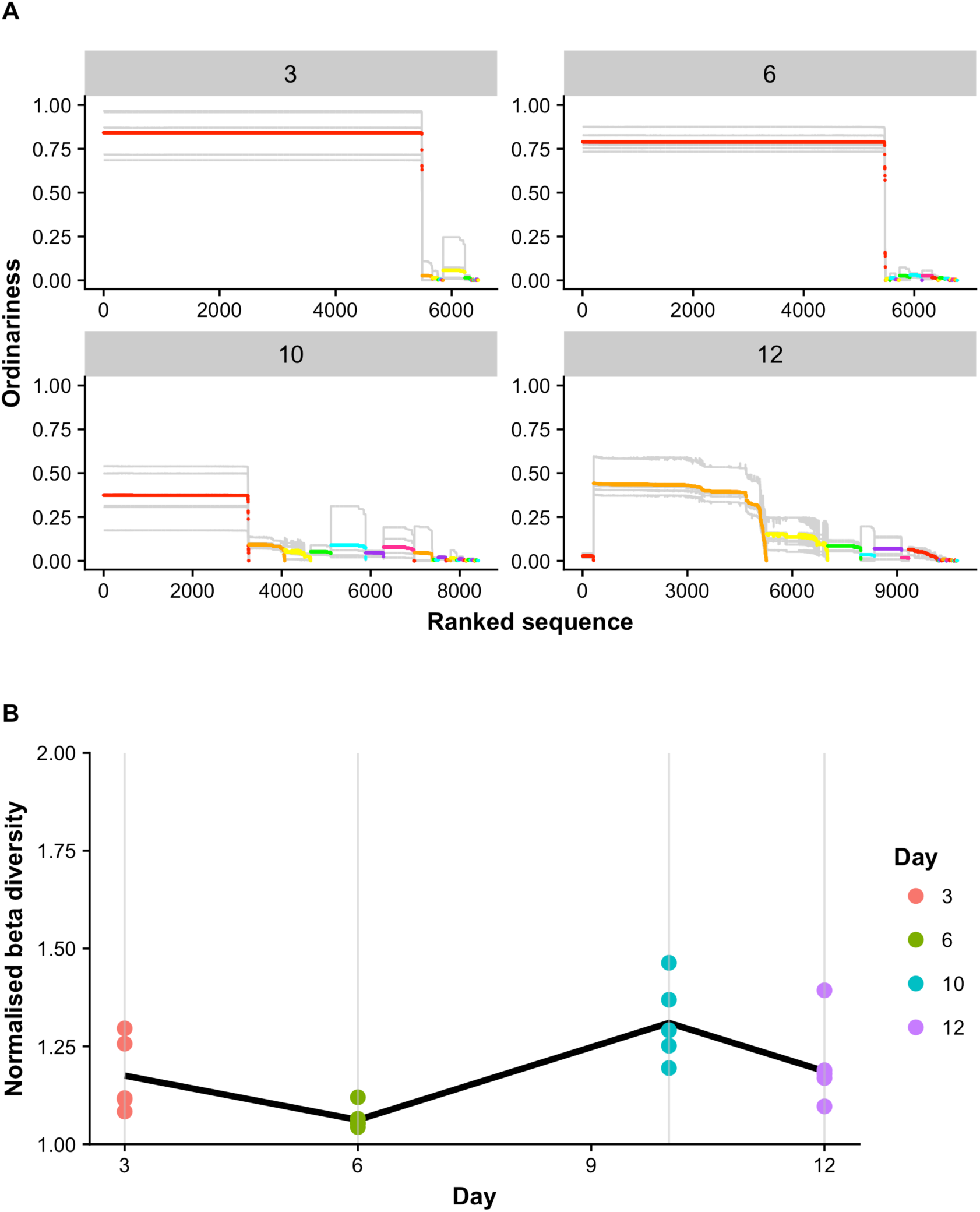
VSGs are expressed in a semi-predicatable order. (A) Expressed VSG sequences were clustered by sequence similarity (Materials and Methods and S4 Appendix) and ordered on the x-axis according to the cluster they were assigned to. The average similarity of each sequence to others from the same population was calculated (‘ordinariness’, a measure of how common that sequence is) and is plotted for each sequence on the y-axis. Grey lines indicate the profiles of individual mice, while coloured lines indicate the average for that particular cluster, coloured according to the cluster. (B) Diversity analysis showing the effective number of distinct VSG profiles found on day 3, 6, 10 and 12 for each mouse (dots) on that day, with the average across the days represented by the solid line.

The sequence similarity data also allowed the analysis of variability between mice using a new measure of population differentiation called normalised beta diversity [30] (Figure 4B). When looking at a single day, beta diversity is the effective number of distinct VSG profiles present on that day, giving information on the differentiation between the animals. This analysis indicates (similar to the VSG mapping data) the greatest beta diversity across individuals is at day 10 (Figure 4B solid line).

Further exploring each time point and variation between mice (Figure 4B dots), we can see that although the mice at day 3 show some distinct VSG profiles (albeit with overexpressed VSGs in individual mice common to all mice, SI Figure 3), at day 6 most mice (except for mouse 6.5) are broadly consistent with respect to which VSGs are present and how common they are. The effective number of VSG profiles increases further on day 10 with maximal divergence between mice at any time point, (Figure 4B, solid line). This value then decreases on day 12 (though mouse 12.2 is distinct), as the mice begin to express similar profiles again. These analyses again indicate that there is stochasticity in the process of VSG expression considered as a progression over 12 days, and there is semi-predictability rather than strict hierarchical progression through VSG expression, as has been described previously [21, 29, 31, 32]

### Mosaic VSG genes are present early in infection

Mosaic genes were identified where BLAST hits for a particular read demonstrated non-overlapping homology to more than one distinct VSG in the reference database. This was commonly seen in the CTerminal domain, where the same N-Terminal domain was in many instances observed with different C-Terminal domains (“3’ donation” in [21]). Using pairwise alignments of all reads that mapped to Tb08.27P2.380, based on the alignment coverage over the gene, donors were filtered based on the region representing the C-Terminal domain (the 3’ region approximating to 30% of the gene shown in Figure 5A). Donors were selected based at least 80% alignment coverage to the CTD. These data show that the reads aligning to Tb08.27P2.380 consists of three subgroups based on their CTD donors, which are derived from either the reference gene Tb08.27P2.380 (43% of all reads), but also from Tb10.v4.0158 (29%) or Tb927.6.5210 (28%). The proportion of the three donor CTDs varies across time points, with the proportion of reads deriving from the donor Tb08.27P2.380 gene decreasing by days 10 and 12 (reducing from 46.55% at day 3 to day 26.19% at day 12, although the number of reads in total aligning to Tb08.27P2.380 is low by days 10 and 12). The frequent nature of this recombination has been observed previously [21]. We detected N-Terminal domain mosaics (within the constraints of our stringent selection criteria) at a much lower frequency (n = 45 over all 20 mice; three sequences at day 3, five at day 6, 13 at day 10 and 23 and day 12 – S3 Table), and in most cases these are single read examples, and so must be treated with some caution (albeit 12 of the putative mosaic reads have coverage of at least 7 full passes, a coverage level at which our analysis – Figure 2D – suggests should effectively remove sequencing-derived error). However, we have two examples where we have more than one read indicating N-Terminal domain mosaicism, with the additional support for one of these sequences that it is only detected in one mouse – given the complex nature of previously identified mosaic N-Terminal domains [5, 21], it is unlikely that identical mosaics would emerge in separate individual infections. Nevertheless, we do also have one putative mosaic sequence that occurs in two separate mice (balbc_6_0/100673/ccs5 and balbc_12_1/30571/ccs9 in mice 6.1 and 12.1, respectively; S3 Table) – this may either represent a gene currently not annotated in the TREU927 genome or be a true mosaic gene that was present in the initial inoculum and has remained at low levels throughout infection. The N-Terminal domain mosaic examples we have detected are mostly relatively simple mosaic genes (e.g. Figure 5B). Although at a low frequency, these are the earliest in trypanosome infections that N-Terminal domain mosaic gene formation has been detected, and the increased frequency over time is consistent with expectations that this process is rarest early in infection.

## Discussion

The results illustrate the power of long read sequencing when applied to expressed allelic diversity – we identified 449 VSGs across 20 individual samples, covering four time points post-infection (3, 6, 10 and 12 days). The identification of the VSGs was achieved by two approaches; mapping reads to a reference VSG database, and clustering read sequences to identify distinct variants – these independent approaches were highly congruent in the number of VSGs and the proportion of reads that were attributed to individual VSGs (Table 2). When compared with previous approaches, such as manual cloning (801 VSG sequences that comprised 93 distinct VSGs or ‘sets’ across 11 mice across 19 days of sampling each [21]) or short read Illumina sequencing (289 VSGs for 4 mice – 3 mice sampled 9 times over 30 days and one mouse sampled 13 times over 105 days), the Pacbio approach gives significantly higher resolution per sample. It must be acknowledged that in the present study the starting volume of infected blood for each sample was higher (200 µl versus 50–100 µl in [23] and approximately 15 µl in [21]), and additionally the inoculum in the current study was significantly greater and not clonal, meaning the study design may predispose to more expressed variants being detectable. The TREU927 clone used was also highly virulent, giving rise to a high parasitaemia early that was maintained for the 12 days of infection – this is not representative of the classical fluctuating profile of less virulent strains (or clones of this strain, e.g. [33]); however, for the purposes of assessing the utility of pacbio this was advantageous. A proportion of the identified VSGs (115/449) derive from single reads in single samples, and therefore a degree of caution must be employed with these variants. However, when the singleton VSGs are removed, we can confidently conclude that we have identified 334 VSGs across our datasets – this ranges from a median of 27 VSGs in day 3 samples to 170 in day 10 samples. Therefore, despite these caveats, we can still conclude that the resolution in terms of diversity is significant for the long read approach, and likely to be of great utility for studies incorporating VSG diversity going forward.

Despite the limitations of the study design, where we have analysed batches of mice at four time points rather than longitudinal surveys of individual mice, our data across 20 mice and four time points are very consistent with a highly reproducible of pattern of VSG expression over time (Figure 3 & Table 2). There was a remarkable degree of consistency in identity of dominant VSGs across independent infections – particularly as the inoculum used was not a single cell or a cloned inoculum (this is very distinct from, for example, *Borrelia*, where Pacbio analysis has indicated very little overlap in expressed antigen diversity across replicates from the same starting inoculum [34]). The data demonstrated a consistent emergence of the two sequentially dominant variants at the beginning and end of the infection period (Tb08.27P2.380 and Tb09.v4.0077), although during the period in between the dominant VSGs there was significant diversity in expressed VSGs that was consistent with an inherent degree of stochasticity in the system. This was reinforced by the application of biodiversity analysis (Figure 4), which illustrated the semi-predictable nature of the variant progression across the mice and timepoints. This chimes with previous work that described the semi-hierarchical expression of VSGs in *T. brucei* [21, 28, 29], and modelling approaches that have also reflected semi-ordered use of the VSG repertoire [31, 32, 35].

When analysing our data set and comparing with that of Hall et al, 2013, who used the same TREU927 strain, we have significant overlap in detected expressed variants. 90% of our reads correspond with a VSG detected in Hall et al. The dominant early VSG is different (corresponding to ‘Set_23’ in the Hall data, Table 2), although the Tb09.v4.0077 which becomes dominant by day 12 was similarly dominant by ∼day 20 in Hall et al; differences are presumably due to the use of either a stabilate with a distinct passage history, or the use of a larger inoculum rather than single trypanosomes (i.e. inoculation of a population from a previous infection presumably expressing the dominant VSG at that particular stage). The dominant VSG in our dataset (Tb08.27P2.380) was annotated as a pseudogene in the reference genome (predicted to be truncated due to insertion of a stop codon); this is not consistent with our data as a dominant early VSG as it would suggest mosaic gene formation providing a dominant early gene – indeed, recent reannotation has classified this gene as intact, which would be more in keeping with early expression favouring intact over pseudogene or mosaic VSGs [5, 21] (although given the 1 × 105 inoculum used in this study, it is feasible that the transfer of Tb08.27P2.380 as the dominant expressed VSG from the donor mouse infection may have given rise to this).

We have identified mosaic genes (classified as reads demonstrated non-overlapping homology to more than one distinct VSG N-Terminal domain in the reference database) earlier in infection than has previously been identified. The rate of mosaic gene formation was very low in our study, mostly either single or very few reads, which probably reflects our timeframe being only 12 days post-infection. The nature of the long read sequencing is highly beneficial in terms of mosaic gene identification; even low frequency expressed genes (within the limitation of the four orders of magnitude of coverage that the read number per sample provides) can be identified with some confidence due to the acquisition of the whole gene sequence – in order to achieve this with short read approaches a reasonable degree of read coverage would be required to identify and confirm putative mosaic genes. This has potential implications for the application of long read sequencing to significantly further our understanding of infection dynamics and the role of mosaic genes as infections progress. This is likely to be important in terms of ability to gain insights into the mechanisms of mosaic gene formation because of consequent increased ability to resolve defects in switching rate (e.g. analysis of DNA recombination gene mutants such as RAD51 that have been implicated in DNA recombination-based VSG switching [15]) – at present it is not known if mosaic gene formation involves a mechanistic switch in terms of pathways; the ability to detect low frequency mosaic gene expression should provide the ability to study this. Additionally, detection of low frequency VSGs would enhance the ability overall to more fully analyse the temporal kinetics of VSG switching – providing an avenue for improved quality of inputs into modelling dynamics of VSG expression. The approach of mapping reads to a defined VSG reference database, and in particular the clustering approach developed in this study, would also make analysing expressed VSGs in the animal trypanosomes, *T. congolense* & *T. vivax*, feasible (one challenge being the lack of conserved 3’ UTR sequence in expressed VSGs to enrich transcripts in both species); this may be particularly enlightening given different structure and content of VSG repertoires recently described between the three genomes [36, 37].

We detected indels consistently when comparing Pacbio transcripts to the reference gene (Figure 2). While these differences may indeed be real, with our protocol we have somewhat limited resolution for conclusively differentiating indels introduced by the trypanosome from those potentially introduced by PCR. However, PCR is unlikely to be the sole cause of the observed mutations, because in the dominant VSG in our dataset (Tb08.27P2.380), which represents 141,822 reads across all 20 samples – therefore, 20 independent PCR reactions – we observe a consistent set of variations from the reference genome sequence (10% mismatches, 1.47% deletions, 1.07% insertions average per base) across all reads – these are consistently present across all reads for this variant, including those reads with high fold coverage (i.e. greater than 10 full passes per read) (Figure 2). These data, across technical and biological replicates, lead us to conclude that these differences were present in most likely the genome copy, but also potentially a distinct BES-resident copy of this VSG that has accumulated mutations distinct to the genome basic copy of the gene, and mutations were not introduced by PCR. One possible explanation for this is that there is very likely a significant (and unknown) divergence in passage history between the sequenced reference genome TREU927 trypanosomes and those used in this experiment. This would be consistent with data from many pathogens of the increased mutability of telomeric/subtelomeric gene families [38]. Previous data indicated accumulation of point mutations in expressed VSGs over time within infections [5, 21], and in our data we saw some support for this process, but the skewed nature of the data distribution limits our ability to conclude increased mutations over time as an important aspect of VSG expression, and it should be noted that a timeframe of 12 days is relatively short and will have limited our resolution – however, our data indicate that application of long read analysis over longer infection timeframes is likely to be a useful means of characterising the nature and role of this mechanism.

However, the multiple mutations that were present across multiple VSG sequences in our data, did enable detailed analysis of the nature of mutations detected in Pacbio sequencing (Figure 2). Ideally, to enable clear differentiation of PCR bias and artefact, errors introduced by Pacbio, and mutations introduced by the trypanosome, unique molecular identifiers (UMIs) would be added prior to PCR amplification (e.g. [22, 39]). While we did not incorporate this step, we can draw some conclusions from analysis of our data. When data for Tb08.27P2.380, which represents 141,822 reads, is analysed across the range of fold coverage per read, it is clear that most of these mutations are removed as the coverage increases (Figure 2) – although notably even at a high number of passes (>10) some introduced mutations remain. This strongly suggests that most of these are errors that are introduced by the Pacbio process, and the proportion we observed across the dataset (4% per base pair) is consistent with that reported in other studies (e.g. [40]). The mutations also directly influenced the ability to predict open reading frames in our data – ORFs only being detected in 11.22% of VSG reads (33,234 of 296,937). Clearly, with these reads being generated from cDNA one would have expected most if not all to have identifiable ORFs. Therefore, these data indicate some of the limitations when using Pacbio, even with data that comprises multiple passes – the introduction of mutations does provide a layer of complexity to the analysis that must be addressed with care. This is particularly pertinent when trying to analyse multiple closely related alleles, as in the case of VSGs – we were able to draw conclusions on the basis of sufficient coverage of a highly expressed dominant allele, combined with the inclusion of multiple biological replicates; without these elements interpretation would have been very difficult without parallel short read sequencing to correct errors introduced by the technology.

A further issue for consideration for the application of long read technologies to the analysis of expressed allelic diversity is the number of reads per sample; with coverage over four orders of magnitude – although significantly greater in resolution than previous manual and laborious methods, this contrasts relatively poorly with the numbers of reads that short read applications deliver (millions) – although it should be noted that with the short read approach many reads will be required to robustly identify full length single variants (in particular to enable differentiation of closely related transcripts, either similar genome-encoded alleles or related lineages of mosaic genes [5, 21]), whereas in theory at least a single pacbio read should provide the ability to robustly identify a particular VSG transcript. While the coverage is being improved with the newer platforms (e.g. the Pacbio Sequel potentially delivers a further tenfold increase in data per run), this may limit resolution in terms of detecting minor variants, for example. We did detect significant expressed diversity, and this is partly explained by our use of a relatively large inoculum, that was not cloned, of a virulent isolate that resulted in high and sustained parasitaemia – therefore, we started with what was probably a relatively diverse population (albeit dominated by expression of Tb08.27P2.380), reflected in the diversity of VSGs detected at day 3 post-infection, which would be significantly lower in the event of a clonal or smaller initial inoculum.

While our data indicate that long read sequencing provides increased resolution in terms of identifying VSG diversity, clearly questions still remain. For example, why the VSG repertoire is so evolved and large? Our data suggest an increased proportion of repertoire is involved, even at early stages, compared to previous studies, which indicates a bigger proportion of the repertoire may be utilised during the lifetime of an infection (which in cattle can be many hundreds of days) than previous data suggests. This is consistent with the data of Mugnier et al [23], where multiple minor variants were observed using an Illumina sequencing approach. However, that study and ours both have limitations, one with relatively few biological replicates (albeit one mouse was followed for ∼120 days) and one that only ventured to 12 days post-infection. Therefore, assessing antigen dynamics in the chronic phase of infections with tools that give significant resolution of expressed antigen diversity will be critical to furthering our understanding of the mechanisms of trypanosome antigenic variation. Key to studying this will be analysing the picture in the truly chronic stages of infection (as was done by Mugnier et al in the context of mouse infections), but particularly doing so in relevant hosts (e.g. cattle [41]) where the total population of trypanosomes in the animal will be potentially 1,000 times greater at peak parasitaemia and where infections may last for 100s of days – this will have a profound influence on the usage of the repertoire (our data, for example, was representative of a total population of approximately 1 × 10^8^ parasites per mouse). Additionally, recent studies indicates *T. brucei* populations inhabit different niches in the mammalian host (e.g. skin and adipose [42, 43]), to the extent that some show evidence of local adaptation with respect to metabolism ([43]) – how this population compartmentalisation interacts with antigenic variation and immunity is likely to be important for parasite maintenance and transmission. Therefore, understanding the dynamics in both the chronic stages of infection and in clinically relevant hosts will potentially provide ideas on the selective pressures that maintain such an elaborate system. Additionally, given the significant advantages described above in terms of identifying low frequency variants (including mosaic VSGs), it may be that a combined long and short read approach is likely to be the optimal way of holistically and accurately identifying expressed VSG diversity; the increased read number of short read technologies in combination with the better resolution of long read technologies would provide significant power to examine the complexity of VSG expression in trypanosomes.

## Materials and Methods

### Trypanosomes and mouse infections

All mice were infected with *Trypanosoma brucei brucei* TREU927, the genome reference strain [19, 44]. A cryostabilate from liquid nitrogen was thawed and inoculated into BALB/c mice in order to amplify a viable in vivo population. Donor mice were euthanased at first peak parasitaemia (approximately 1 × 10^7^ trypanosomes/ml), and blood extracted. Trypanosomes were counted in triplicate under an improved Neubauer haemocytometer, diluted to inocula of 1 × 10^5^ trypanosomes in 200 µl Carter’s Balanced Salt Solution, which were then inoculated via the intraperitoneal route into 20 recipient BALB/c mice. Mice were maintained for 12 days post-infection, and groups of 5 mice were euthanased at 3, 6, 10 and 12 days post-infection. Parasitaemia was monitored daily by venesection of the lateral tail veins using the rapid matching technique [45], and was counted in triplicate under an improved Neubauer haemocytometer on the sampling days. Care and maintenance of animals complied with University regulations and the Animals (Scientific Procedures) Act (1986; revised 2013).

### RNA extraction, cDNA generation & PCR amplification of VSG transcripts

At each sampling day, RNA was extracted from 200 µl infected blood using the Qiagen RNeasy kit (Qiagen), according to the manufacturer’s instructions. Approximately 1 µg RNA was treated with DNase Turbo (Ambion), according to manufacturer’s instructions, and cDNA was generated as in Hall et al, 2013 [21], including a column purification step on generated cDNA using the PCR Purification kit, according to the manufacturer’s instructions (Qiagen). VSG transcripts were enriched by carrying out PCR with proof reading Herculase II Fusion polymerase (Agilent) on the cDNA template with oligonucleotide primers specific to the *T. brucei* spliced leader sequence (TbSL) and a reverse primers complementary to a 13 base pair conserved region in VSG 3’ untranslated regions (3UTR); primer sequences and PCR conditions were as previously described [21, 23]. A subset of PCR transcripts was subjected to cloning and sequencing; PCR products were ligated into pGEMT-Easy vectors, transfected into One Shot TOP10 cells, bacteria were grown up and cloned under suitable antibiotic selection (all using the TOPO cloning kit, Invitrogen), and plasmid DNA extracted using a Miniprep kit (Qiagen); these procedures were all carried out according to manufacturer’s instructions. Extracted plasmid DNA of appropriate concentration was sent for sequencing (Eurofins MWG).

### Pacbio sequencing

1 µg of PCR amplicon template as measured by Nanodrop (ThermoScientific) and Bioanalyser (Agilent) was submitted to the Centre for Genomic Research, University of Liverpool for sequencing using the Pacbio RSII platform (Pacific Biosciences). DNA was purified with 1x cleaned Ampure beads (Agencourt) and the quantity and quality was assessed using Nanodrop and Qubit assay. Fragment Analyser (using a high sensitivity genomic kit) was used to determine the average size of the DNA and the extent of degradation. DNA was treated with Exonuclease V11 at 37 **°**C for 15 minutes. The ends of the DNA were repaired as described by the Pacific Biosciences protocol. Samples were incubated for 20 minutes at 37 **°**C with damage repair mix supplied in the SMRTbell library kit (Pacific Biosciences). This was followed by a 5 minute incubation at 25 **°**C with end repair mix. DNA was cleaned using 0.5x Ampure beads and 70% ethanol washes. DNA was ligated to adapter overnight at 25 **°**C. Ligation was terminated by incubation at 65**°**C for 10 minutes followed by exonuclease treatment for 1 hour at 37**°**C. The SMRTbell libraries were purified with 0.5x Ampure beads. The quantity of library and therefore the recovery was determined by Qubit assay and the average fragment size determined by Fragment Analyser. SMRTbell libraries were then annealed to the sequencing primer at values predetermined by the Binding Calculator (Pacific Biosciences) and a complex made with the DNA Polymerase (P4/C2chemistry). The complex was bound to Magbeads and this was used to set up 3 SMRT cells for sequencing. Sequencing was done using 180 minute movie times. Data (raw sequencing files) is available through Gene Expression Omnibus (https://www.ncbi.nlm.nih.gov/geo/ – accession number GSE114843).

### Pacbio Sequencing analysis

#### Raw data processing

Pacbio raw data was initially processed using the Pacbio SMRT analysis protocol (v2.3), to convert the data into a fasta file using the following parameter selections: minimum 1 full pass, minimum predicted accuracy of 90%. Based on the read length distribution, a range of 1400–2000bp was used to filter the sequenced reads for downstream VSG analysis.

#### VSG read analysis

Preliminary VSG variant distribution was determined by locally aligning the reads to TREU927 reference transcripts. We generated a local database of TREU927 VSGs, by downloading all transcripts annotated as ‘VSG’ from the most recent version of the TREU927 genome (v26) on www.tritrypdb.org [25]. This resulted in a reference library of 1,557 VSG sequences (including all gene fragments and pseudogenes annotated as VSG – available through GEO accession number GSE114843). This reference set was used to set up a local BLAST+ [46] database to create files of curated protein and nucleotide sequences. Reads were blasted (BLASTn) against the reference VSG database to identify the donor gene. A minimum alignment coverage of 60% or above to the sequence read was used to identify the dominant donor transcript, and to generate a variant distribution chart for each sequenced sample. We also locally aligned the sequenced VSG reads to a blast database of 515 previously identified cloned reads from *T. brucei* TREU927 infections [5, 21](available through GEO accession number GSE114843)

#### Mosaic gene identification

We reasoned that putative mosaic genes could be identified as PacBio sequences with partial, nonoverlapping alignments to multiple VSG genes. We therefore undertook full pairwise alignment using local blast of the 296,937 reads that align to VSGs at a 60% identity threshold post size-selection filtering (see above) against the curated VSG database described above. This resulted in all possible donors and their alignment regions for any specific read being identified.

To distinguish mosaics where the same NTD region occurs with multiple CTD regions (which happens frequently) from mosaics with multiple NTD donors (which are rare), we plotted the number of donor alignments per nucleotide across all reads, with read (VSG) scaled to 100 to enable comparison across multiple variant lengths – this also enabled analysis of the number of donor VSGs across the scaled VSG representatives of our 296,937 VSG dataset. A distinctive increase in the number of alignments was observed at approximately 75% of the sequence length (consistent with the start of the CTD), and we conservatively defined the NTD region of each sequence as the first 70% of its nucleotides (see Figure 5A). This allowed us to define approximate NTD regions of all sequences, including those without ORF.

**Figure 5.**
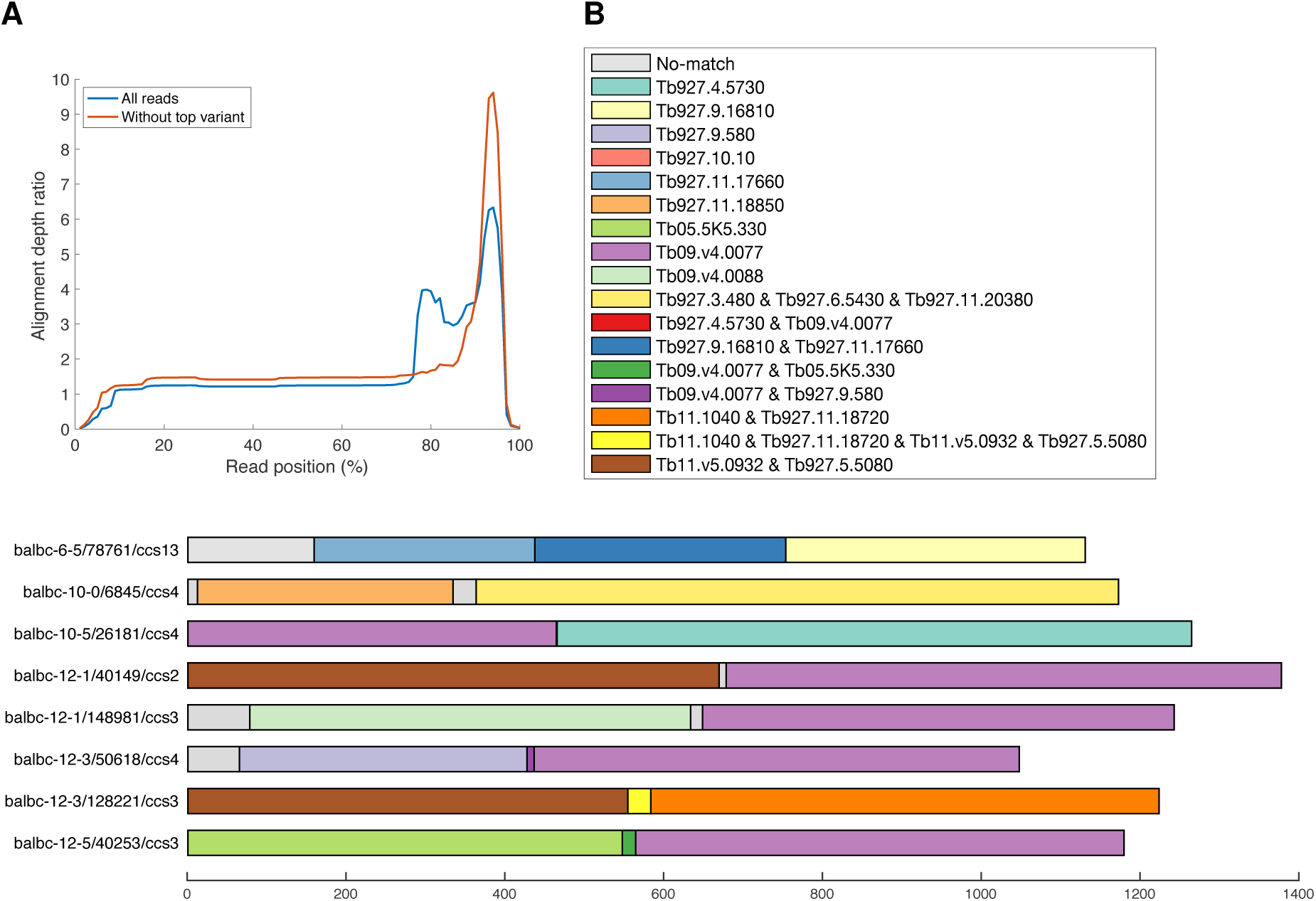
Mosaic gene identification. (A) Average number of donor VSGs (y-axis) per position across scaled VSG reads (gene length scaled to 100 (x-axis); data derived from 296,937 VSG amplicons) – blue line shows data for all reads, red line shows data for all data without reads that map to Tb08.27P2.380. (B) Example mosaic gene candidates, represented by read identifier, and displayed to scale from 5’ to 3’ (N-terminal domain only), with colour of each segment indicating most likely donor VSG from the TREU927 VSG repertoire (VSG gene shown in key; where relevant sequences are identical for more than one potential reference gene donor; all possibilities are shown in key).

Pairwise alignments were then filtered based on the criteria that the start of the alignment should be within the NTD region, and the remaining alignments were used to generate parameters for each read, including NTD length, alignment coverage start and stop sites, NTD alignment coverage percentage, number of donor sequences, alignment coverage of the longest donors, and difference (expressed as percentage non-identity) between the total aligned region and the top donor alignment. Most sequenced NTD regions resulted in full length or partial match to the known VSG database. In order to confidently identify putative mosaic genes, data were further filtered based upon the following criteria; (1) number of donor VSGs is more than one, (2) alignment coverage of the largest donor is less than 80%, and (3) the difference in alignment regions between donors is greater than 10% of the sequence. The remaining sequences were then inspected manually to select mosaic genes.

#### Software

All scripts for raw data processing, VSG read analysis and mosaic gene identification–are available through GitHub (https://github.com/siddharthjayaraman/longread-application).

#### Clustering analysis

We used similarity-based clustering to identify VSG clusters among the sequences. Since, as described above, PacBio reads are prone to introduction of insertion and deletion indels, to reduce the impact of these errors on the quantification of variants detected, we proceeded with only those reads which generated an ORF longer than 400 amino acid residues. We pool all of these sequences from each day and each mouse. We used Clustal Omega to calculate genetic distances between each pair of sequences [47] and clustered sequences using a 6% threshold (employed in many clustering algorithms, e.g. UClust) for intra-cluster dissimilarity. To resolve the problems surrounding cluster identification a novel dynamically resizing clustering algorithm was implemented (full details are given in S4 Appendix).

#### Diversity analysis

For a de-novo approach to quantify the observed variation in VSG variants in our samples over time we used novel diversity metrics [30], that have been developed in theoretical ecology in order to measure biodiversity across scales. We regard VSGs as the ‘ecological species’ in this setting, so in the very simplest case biodiversity would simply be how many species or VSGs we observe. The measures go a step further than this and weight for the relative abundance of the VSGs via a parameter q (in the main text we use q = 1, for other values of q and a discussion of this see S4 Appendix). In addition, the measures account for the similarity of the sequences in such a way that if two sequences differed in only one base pair they would be essentially regarded as the same “species” or VSG, as they would have close to 100% similarity.

Within the diversity framework, normalised beta diversity [30] quantifies population differentiation. We consider the VSGs from all mice on a given day as our population (the *metacommunity* level) and normalised beta diversity measures the number of distinct mouse (the subcommunity level) VSG *profiles* that are present on that day (see S4 Appendix, S5 & S6 Figures).

## Acknowledgements

This work was funded by a Royal Society University Research Fellowship (UF090083 & UF140610) to LM, and a Royal Society grant (RG110378) to LM and CHF. TM and LM were funded by a Roslin sequencing grant (BBSRC). The Roslin Institute is supported by the BBSRC (BB/J004227/1, BB/J004235/1 & BBS/E/D/20002173). RR and CC are supported by the BBSRC (BB/P004202/1).

## Supporting Information

S1 Table.

Amplicon read count per sample for each identified VSG transcript (xlsx file).

**S2 Figure.**
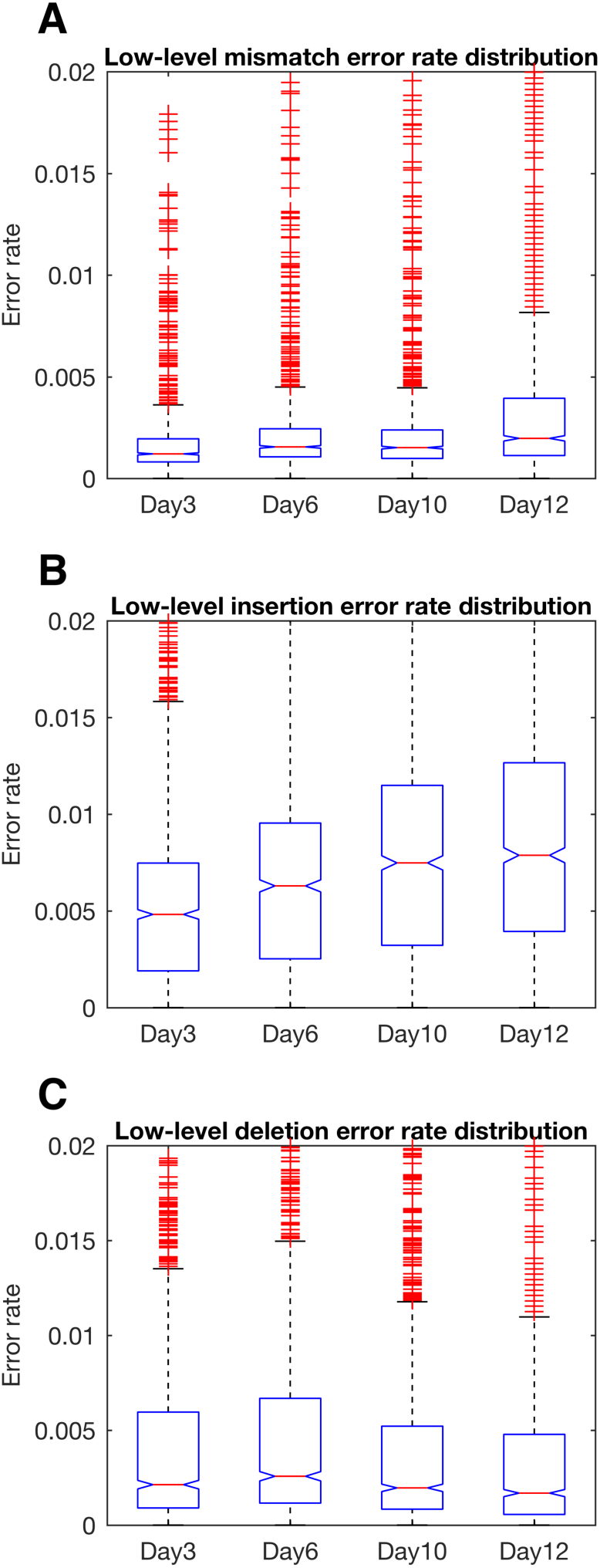
Error (mutation) rate distribution over time (day 3, 6, 10 and 12 post-infection) for reads aligning to VSG Tb08.27P2.380 for (A) mismatches, (B) insertions and (C) deletions; each defined as differences relative to the reference genome sequence of Tb08.27P2.380. For each mutation class and timepoint, the boxplot showns median values and 25th and 75th percentiles, whiskers extend to data extremes, and data outliers are plotted individually (red plus symbols).

**S3 Table.** Putative VSG mosaic transcripts.

| Day3 | Day6 | Day10 | Day12 |
| --- | --- | --- | --- |
| balbc_3_4/151297/ccs3 (3.5) | balbc_6_0/100673/ccs5 (6.1) | balbc_10_0/102388/ccs3 (10.1) | balbc_12_1/114076/ccs2 (12.1) |
| balbc_3_4/156604/ccs3 (3.5) | balbc_6_1/88751/ccs2 (6.2) | balbc_10_0/118718/ccs6 (10.1) | balbc_12_1/146680/ccs3 (12.1) |
| balbc_3_4/75350/ccs6 (3.5) | balbc_6_2/21985/ccs8 (6.2) | balbc_10_0/128573/ccs2 (10.1) | balbc_12_1/148981/ccs3 (12.1) |
|  | balbc_6_5/151460/ccs7 (6.5) | balbc_10_0/48011/ccs2 (10.1) | balbc_12_1/30571/ccs9 (12.1) |
|  | balbc_6_5/78761/ccs13 (6.5) | balbc_10_0/6845/ccs4 (10.1) | balbc_12_1/40149/ccs2 (12.1) |
|  | balbc_6_5/86021/ccs7 (6.5) | balbc_10_2/138490/ccs5 (10.3) | balbc_12_1/48081/ccs4 (12.1) |
|  |  | balbc_10_4/106314/ccs2 (10.4) | balbc_12_1/9710/ccs9 (12.1) |
|  |  | balbc_10_4/13775/ccs2 (10.4) | balbc_12_1/97157/ccs2 (12.1) |
|  |  | balbc_10_4/82822/ccs2 (10.4) | balbc_12_2/126314/ccs8 (12.2) |
|  |  | balbc_10_5/110598/ccs2 (10.5) | balbc_12_2/90072/ccs7 (12.2) |
|  |  | balbc_10_5/125683/ccs15 (10.5) | balbc_12_3/102971/ccs2 (12.3) |
|  |  | balbc_10_5/14682/ccs9 (10.5) | balbc_12_3/105707/ccs7 (12.3) |
|  |  | balbc_10_5/26181/ccs4 (10.5) | balbc_12_3/128221/ccs3 (12.3) |
|  |  |  | balbc_12_3/156428/ccs2 (12.3) |
|  |  |  | balbc_12_3/40832/ccs2 (12.3) |
|  |  |  | balbc_12_3/50618/ccs4 (12.3) |
|  |  |  | balbc_12_3/67024/ccs3 (12.3) |
|  |  |  | balbc_12_3/75428/ccs8 (12.3) |
|  |  |  | balbc_12_3/83905/ccs2 (12.3) |
|  |  |  | balbc_12_5/135299/ccs5 (12.5) |
|  |  |  | balbc_12_5/145884/ccs2 (12.5) |
|  |  |  | balbc_12_5/40253/ccs3 (12.5) |
|  |  |  | balbc_12_5/46204/ccs6 (12.5) |

\*The coloured cells represent identical sequences

## S5 Appendix. Clustering algorithm and Diversity analysis detailed methods

### Clustering algorithm

Performing a non-dynamics clustering analysis, using a fixed standard 6% threshold for intra-cluster dissimilarity we observed that most sequences fell into clusters, but there was a large number (∼6000) of unclustered reads many of which were only just outside the 6% threshold for a cluster and so should have been assigned to that cluster. To address this a dynamic clustering algorithm was designed which would allow the cluster threshold to grow for each cluster independently allowing us to correctly deal with the unclustered reads. The algorithm proceeded as follows:

1. We used Clustal Omega to calculate genetic distances between each pair of sequences [45] and converted these into similarities using a 6% threshold for dissimilarity. All sequence pairs below the threshold for dissimilarity were given a similarity of 1 and those above given 0. This strict cutoff was chosen to allow our method to be comparable to traditional clustering approaches.
2. Centroid sequences that would start off each new cluster were chosen using the core measure from the diversity framework, ordinariness [46]. Initial ordinariness O_1i_ of sequence i is the average similarity of any sequence to sequence i, or here the number of sequences within 6% sequence dissimilarity of sequence i. The sequence with the highest ordinariness was chosen as the centroid of the first cluster, and all sequences within 6% genetic distance were included in that cluster.
3. Ordinarinesses, O_ci_, were then recalculated for cluster c (c = 2, 3 …) from all of the remaining sequences not currently allocated to a cluster and the method for identifying a new centroid (step 2) was repeated.
4. From the third cluster onwards (c > 2), however, any sequence with a higher initial ordinariness O1i than the previous cluster centroid must have derived that ordinariness from similarity to an earlier cluster (as otherwise it would have been selected earlier), and is therefore assigned to its closest cluster rather than forming a new cluster. The cluster threshold for that closest cluster is then re-calculated to accommodate the new sequence, and any other sequences that now fall within the new sequence dissimilarity threshold for that cluster are added to it.
5. The clustering algorithm repeats until no more sequences are present or the cluster sizes grow too large and begin to overlap, leaving residual unique (unclustered) sequences.

### Diversity analysis

For the diversity analysis we need to define the *metacommunity* which is the VSG profile on a given day when we pool the data from all the mice from that day (SI Figure 1A). The *subcommunity* is then the VSG profile for a particular mouse on that day. So each metacommunity (day) is made up of 5 subcommunities (mice). By *VSG profile* we mean the distribution of VSGs as illustrated by the pie charts in SI Figure 1A.

When looking at how different each mouse’s VSG profile is within each day one might start by asking: of all the VSGs seen on that day how many are seen in mouse i from that day? If each mouse has half of the possible VSGs from that day then we say there are effectively 2 distinct VSG profiles on that day because this is the minimum number of distinct VSG profiles we would need to account for all the diversity we see, because any given profile only contains half of that day’s diversity. This number of distinct profiles is referred to as *normalised beta diversity* for q = 0, and is calculated for each mouse on a given day (see SI Figure 1B). The parameter q weighs the extent to which the relative proportion of each VSG is taken into account when we assess normalised beta diversity. When q = 0 we ignore the relative proportions of each VSG, we only consider if each VSG is present or not. As q increases we no longer simply care about how many of the VSGs are present in each mouse, but also how faithfully the mouse (subcommunity) preserves the proportions of each VSGs observed across all the mice from that day (metacommunity). Unless the VSGs are evenly distributed across each mouse we find that increasing q results in an increase in the normalised beta diversity as more VSG profiles are needed to make up for the unevenness in the proportions of VSGs.

We see that for our dataset normalised beta diversity, or the effective number of distinct VSG profiles is close to 1 for q = 0 on each day (SI Figure 2B). This is because each mouse on a given day expresses at least some of almost every VSG present on that day, so there is only one VSG profile when we ignore the role of relative abundances of the VSGs. However, as q increases, and we start to care about accuracy of the abundances in the profiles and we are able to detect that some of the mice become increasingly divergent from the rest (e.g. mouse 3.4 and 3.5 in day 3), because they have some VSGs that are more common when compared to the rest of the mice in that day. This is contrast to day 6 where most mice (except for mouse 6.5) broadly agree on not only which VSGs are present (q = 0), but also on how common those VSGs are on that day (shown by the closeness of the lines in SI Figure 2B, day 6 for q>0). Day 10 has the most distinct VSG profiles, with a lot of variability between the mice, while on day 12 the mice begin to express similar VSG profiles again (though mouse 12.2 is distinct).

These results are further supported by additional clustering analysis. We can apply the clustering algorithm to each mouse individually (SI Figure 2). Looking at day 3 we see that the orange and yellow clusters only appear in 3 of the mice on day 3 (mice 3.1, 3.4, 3.5), highlighting the diversity between the mice that we saw from the normalised beta diversity analysis. Similarly, the day 10 mice show a lot of variation in clusters of VSGs they express, whereas on day 12 the clustering is broadly the same for each mouse.

---

**S5 Figure.**
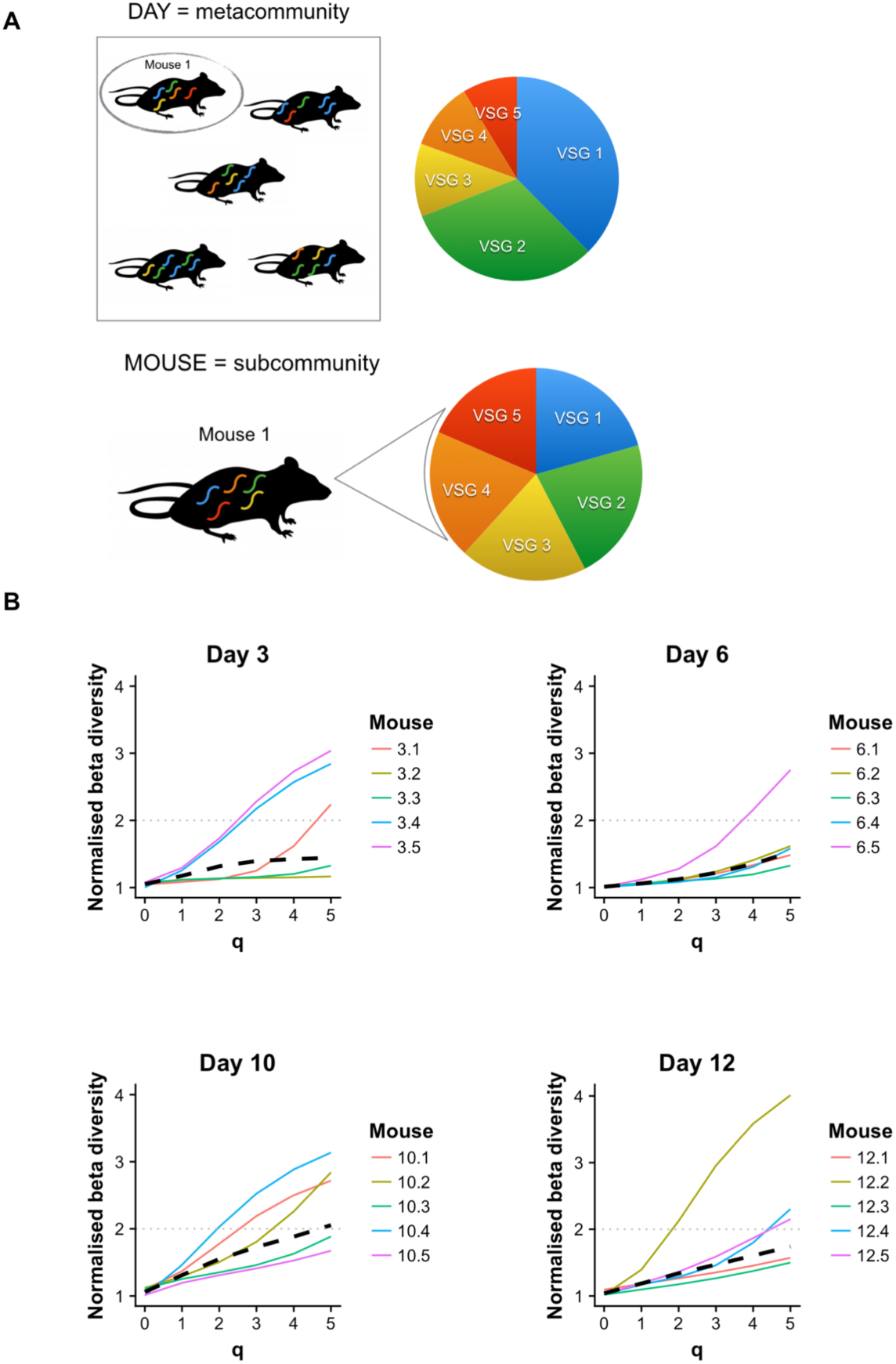
Diversity analysis (A) Structuring of the data for diversity analysis. The combined VSG profile from all mice on a given day form the metacommunity, which is the unit of analysis; the VSG profile from each individual mouse form a single subcommunity of reads within that metacommunity. So each metacommunity (day) is made up of 5 subcommunities (mice). (B) Normalised beta diversity analysis for varying weightings (q) of VSG proportional abundance. The y-axis shows the effective number of distinct VSG profiles found on a given day seen from the perspective of each mouse (coloured lines) on that day, with the average across the day given by the dashed line. The x-axis indicates how much relative proportions of VSGs rather than just the presence-absence of the VSG is weighted in the assessment of diversity. When q = 0 only the presence or absence of the VSG is considered when comparing an individual mouse’s VSG profile to the profile obtained from pooling all the mice from that day. For large q, we compare not only the presence and absence of VSGs but also their relative proportions. The larger the value of q the less importance is placed on rare VSGs in a profile. The more a mouse differs from the pooled data the higher the value of normed beta diversity.

**S6 Figure.**
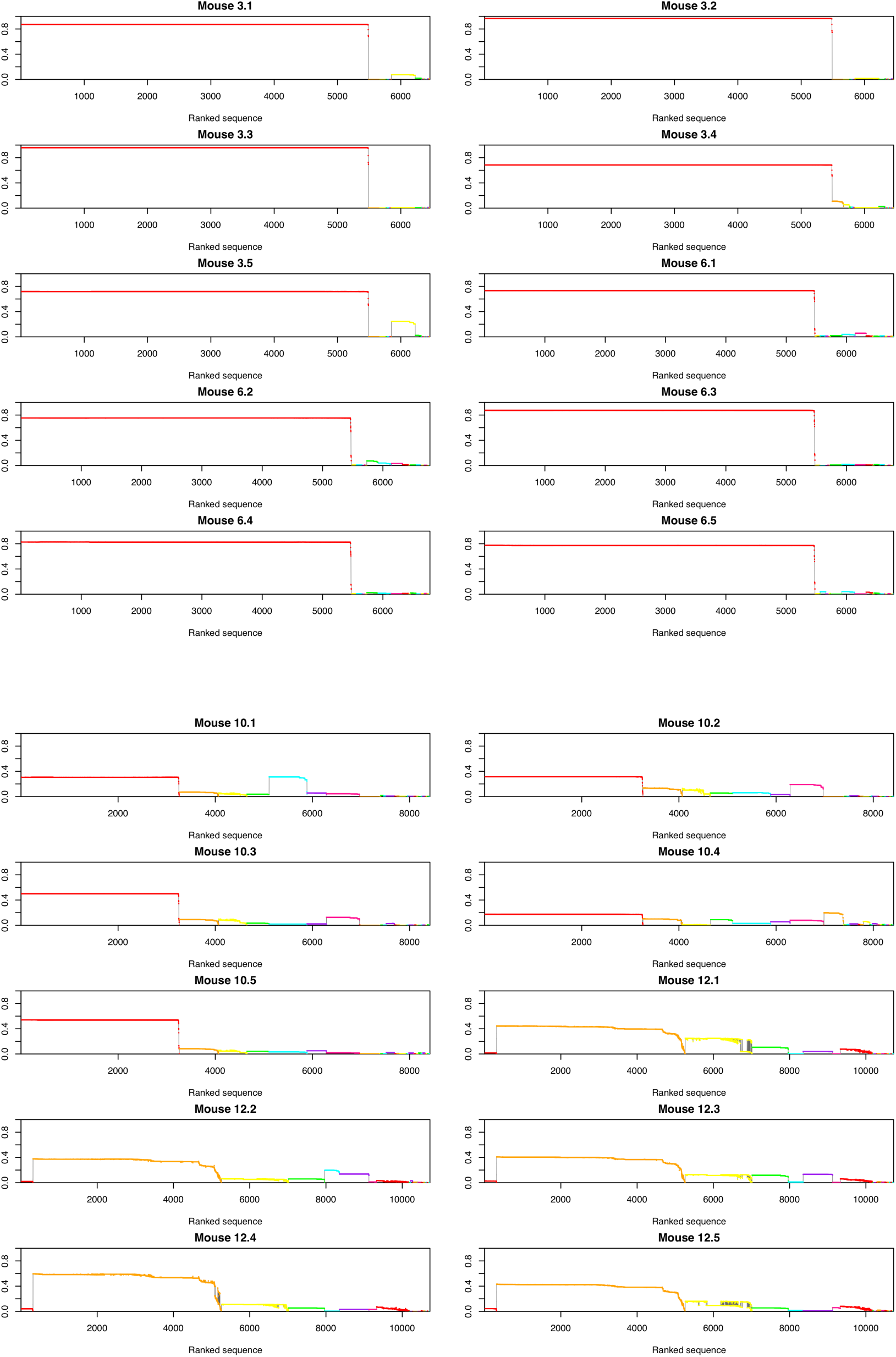
Clustering analysis of reads from each mouse. The y-axis indicates how common the cluster is in that mouse and the x-axis indicates how many sequences fall within that cluster. Clusters are colour coded such that a red cluster in mouse 3.1 is defined by the same centroid and clustering threshold as the red cluster in mouse 10.5 etc.

